# Pazopanib induces dramatic but transient contraction of myeloid suppression compartment in favor of adaptive immunity

**DOI:** 10.1101/2020.05.01.071613

**Authors:** Darawan Rinchai, Elena Verzoni, Veronica Huber, Agata Cova, Paola Squarcina, Loris De Cecco, Filippo de Braud, Raffaele Ratta, Matteo Dugo, Luca Lalli, Viviana Vallacchi, Monica Rodolfo, Jessica Roelands, Chiara Castelli, Damien Chaussabel, Giuseppe Procopio, Davide Bedognetti, Licia Rivoltini

## Abstract

Anti-angiogenic tyrosine-kinase inhibitors (TKIs) and immune checkpoint blockade (ICB) constitute the backbone of metastatic renal cell carcinoma (mRCC) treatment. The development of the optimal combinatorial or sequential approach is hindered by the lack of comprehensive data regarding TKI-induced immunomodulation and its kinetics. Through the use of orthogonal transcriptomic and phenotyping platforms combined with functional analytic pipelines, we demonstrated that the anti-angiogenic TKI pazopanib induces a dramatic and coherent reshaping of systemic immunity in mRCC patients, downsizing the myeloid-derived suppressor cell (MDSC) compartment in favor of a strong enhancement of cytotoxic T and Natural Killer (NK) cell effector functions. The intratumoral expression level of a MDSC signature here generated was strongly associated with poor prognosis in mRCC patients. The marked but transient nature of this immunomodulation, peaking at the 3^rd^ month of treatment, provides the rationale for the use of TKIs as a preconditioning strategy to improve the efficacy of ICB.

## Introduction

The off-target activity of cancer treatments on systemic immunity is a well-known process, deemed to play an active role in disease control [1]. In the course of conventional therapies including chemotherapy or radiotherapy, a reshaping of spontaneous immune response toward a boost of tumor-specific T cells might be required to achieve significant clinical benefit [2]. Such a reinvigoration of adaptive T cell immunity stems from a network of effects triggered by tumor cell death, amplified by additional mechanisms involving multiple immune pathways depending on the drug’s mechanism of action [3].

The immunomodulatory properties of cancer therapies are recently gaining attention in view of potential combinations with immunotherapy such as immune checkpoint blockade (ICB). The nowadays consolidated evidence that ICB mediates effective tumor control only in a minority of patients and in selected malignancies, points to the use of drug combinations as a strategy to increment ICB effectiveness [4]. Promising results showing increased efficacy of PD-1 blockers combined with chemotherapy in non-small cell lung cancer (NSCLC) [5] and breast cancer [6, 7], along with multiple clinical trials ongoing in different tumor types, suggest that combinatorial approaches hold promise to become the potential gold standard of treatment in different settings.

However, the effects of cancer therapies on the multiple components of tumor immunity might be complex and variegated and need to be carefully considered when combination strategies based on desired synergies are designed. Antineoplastic treatments might directly potentiate tumor immunogenicity by broadening antigenic repertoire or favoring antigen presentation that boost T cell priming [5]; or they can act indirectly by reducing myeloid-derived suppressor cell (MDSC)-mediated immunosuppression as a beneficial outcome of their common myelotoxicity [8]. Conversely, according to *in vitro* and/or *in vivo* experimental studies, anti-proliferative therapeutic strategies, particularly those based on the inhibition of multiple tyrosine kinases, might affect T cell proliferation and function as well, thus potentially interfering with the protective activity of adaptive immunity [9–12]. Hence, gaining detailed information on the type and kinetics of the immunomodulating properties of anticancer drugs would be essential to maximize clinical efficacy when diverse therapeutic strategies are combined with immunotherapy.

Given the complexity of the human immune system and its dynamic nature, immunomonitoring approaches are moving toward multiplex and “omics” strategies, with first results emerging in autoimmunity and viral infections [13, 14]. Transcriptomic analysis of peripheral blood [15, 16] for instance, has been extensively used to dissect mechanisms of action of vaccination against infectious diseases [17], to elucidate pathogenic mechanisms of different immunologic disorders [18, 19], and to identify perturbations associated with different viral [20], parasitic [21], or bacterial infections [18, 22]. However, such an approach remains relatively unexplored in the context of cancer therapy, including immunotherapy [23]. Pioneering studies in cancer patients treated with interleukin (IL)-2 have contributed to the characterization of systemic changes induced by this cytokine [24–26]. More recently, peripheral blood transcriptomic analysis has been used to identify signatures associated with responsiveness to anti-CTLA4 [27], and to describe changes differentially associated with CTLA4 and combined CTLA4/PD-1 blockade [23, 28]. To the best of our knowledge, no peripheral blood transcriptomic studies have been performed so far in solid cancer patients receiving tyrosine kinase inhibitors (TKI) or in renal cell carcinoma (RCC) patients treated with any other drugs. According to first-line randomized 3 metastatic RCC trials using PD-1 blockades, up to 50-60% of patients do not respond to such treatment [29–31]. Recently, a retrospective analysis of the CheckMate 025 phase III trial demonstrated a remarkable increased overall survival (HR, 0.60; 95% CI,.0.42–0.84) in patients previously treated with first-line pazopanib [32].

In the present work, we applied an integrative analysis encompassing transcriptional profiling (leucocyte subtype abundance estimation, functional characterization by pathway analysis and modular repertoire analysis) and multiplex flow cytometry, to comprehensively capture the immunomodulation mediated by anti-angiogenic treatment in mRCC patients. RCC was specifically chosen for its potent immunosuppressive properties linked to hypoxia/VHL-related oncogenic pathways that lead to the secretion of proangiogenic factors known to mediate the blunting of adaptive antitumor immunity [33]. Anti-angiogenic drugs interfering with one or multiple factors of the VEGF, PDGF, HGF and ANG2 cascade, are endowed with the intrinsic ability to overcome VEGF-mediated immunosuppression [34, 35]. Here, we investigated the entity and the kinetics of immunomodulation mediated in mRCC patients by pazopanib, a TKI anti-angiogenic drug included in the standard care of this disease [36–38].

By performing a matched analysis of transcriptional and phenotypic profiling in peripheral blood mononuclear cells (PBMCs) obtained before and at different time points during therapy, we demonstrated, for the first time, that pazopanib mediates a potent but transient reprogramming of systemic immunity, resulting in an enhanced T and NK cytotoxic response accompanied by an attenuation of the myeloid suppressor compartment. Our study shows that monitoring systemic immunity by transcriptomics may help designing drug-tailored combination strategies aimed at maximizing clinical efficacy thanks to the timely engagement of a full-fledged immune response.

## Results

### Integrative transcriptional analysis reveals the immune modulatory properties of pazopanib

The study has been conducted on clear cell mRCC patients treated with first line pazopanib, whose PBMCs were obtained from blood withdrawn at baseline, 3- and 6-months during treatment. Transcriptional profiling was analyzed using integrative and complementary pipelines.

We first wanted to explore the molecular heterogeneity of the sample set through principal component analysis (PCA) based on the whole transcriptomic profile (12,913 genes) (**Figure 1A**). The first 3 major PCs accounted for 20.7% (PC1), 10.7% (PC2), and 8.0% (PC3) of the variability observed for these conditions. These three-dimensional plots showed the distribution of individual patient samples for each time point with no outlier sample **Figure 1A**. In general, a certain degree of separation according to different time points was observed.

**Figure 1:**
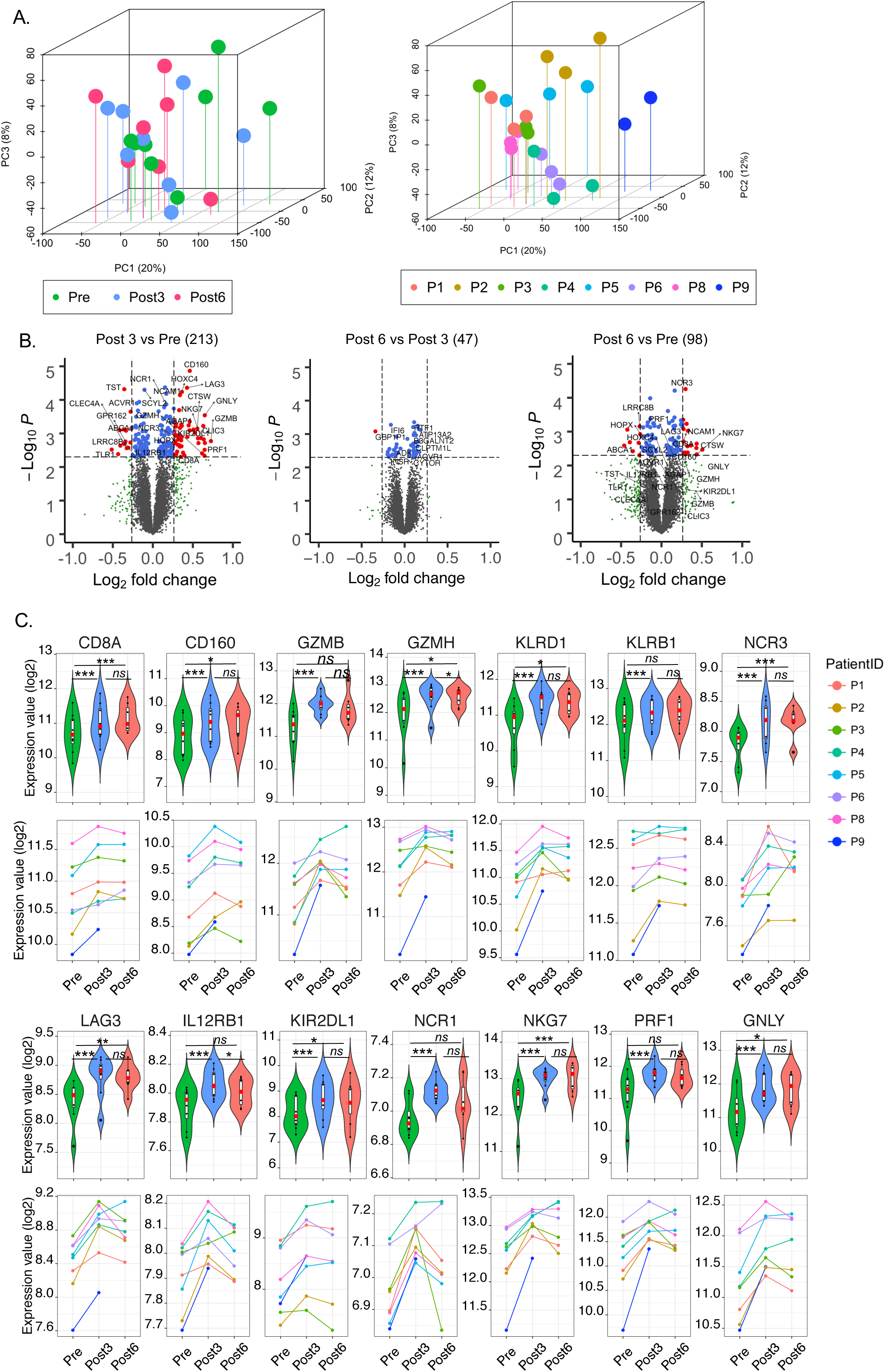
Transcriptional analysis of PBMCs from mRCC patients treated with pazopanib. (A) Principle component analysis (PCA) of all patient samples color-coded by time of treatment (left) and individual patient (right). (B) Volcano plots of differentially expressed genes between pre- and post-treatment (*Post 3* vs *Pre, Post 6* vs *Post 3* and *Post 6* vs *Pre*); the horizontal line represent cut off at *p* <*0.005* and vertical lines represent fold-change > 1.2 (right) or < −1.2 (left). (C) Violin and line plots of selected genes. The paired *t-test* was used for comparison of the expression levels of each gene between patient groups.

We then performed differential expression analysis between post-treatment versus pre-treatment samples. Two hundred and thirteen transcripts were significantly different between 3 months post-treatment (*Post 3*) and pre-treatment (*Pre*), 47 transcripts between 6 months and 3 months post-treatment (*Post 6 vs Post 3*), and 98 transcripts between *Post 6* and *Pre*. Volcano plots showing log2 fold-change (log2FC) and *p-value (paired t-test)* on differentially expressed genes are shown in **Figure 1B**, and **Supplementary Table 1.** Strikingly, among the top 40 genes ranked according to the log2FC, the large majority was associated with cytotoxic functions and interferon signaling (**Table 1**). Representative transcripts related to T and NK cells cytotoxic functions and T cells activation (e.g., *CD8A, CD160, GZMB, GZMH, KLRD1, KLRB1, NRC3, LAG3*, IL12RB1, KIR2DL1, *NCR1, NKG7, PRF1* and *GNLY)* are represented in **Figure 1C**. The over-expression of such transcripts was attenuated at 6 months after treatment. The common transcripts significantly up-regulated at both *Post 3* and *Post 6 (N=16)* as compared to pre-treatment include CD8A, *CTSW, NCAM1 (CD56), NCR3, NKG7*, consistent with a persistence but attenuated NK and T cell response [39] (**Figure 1B**, and **Supplementary Table 1**).

**Table 1:**
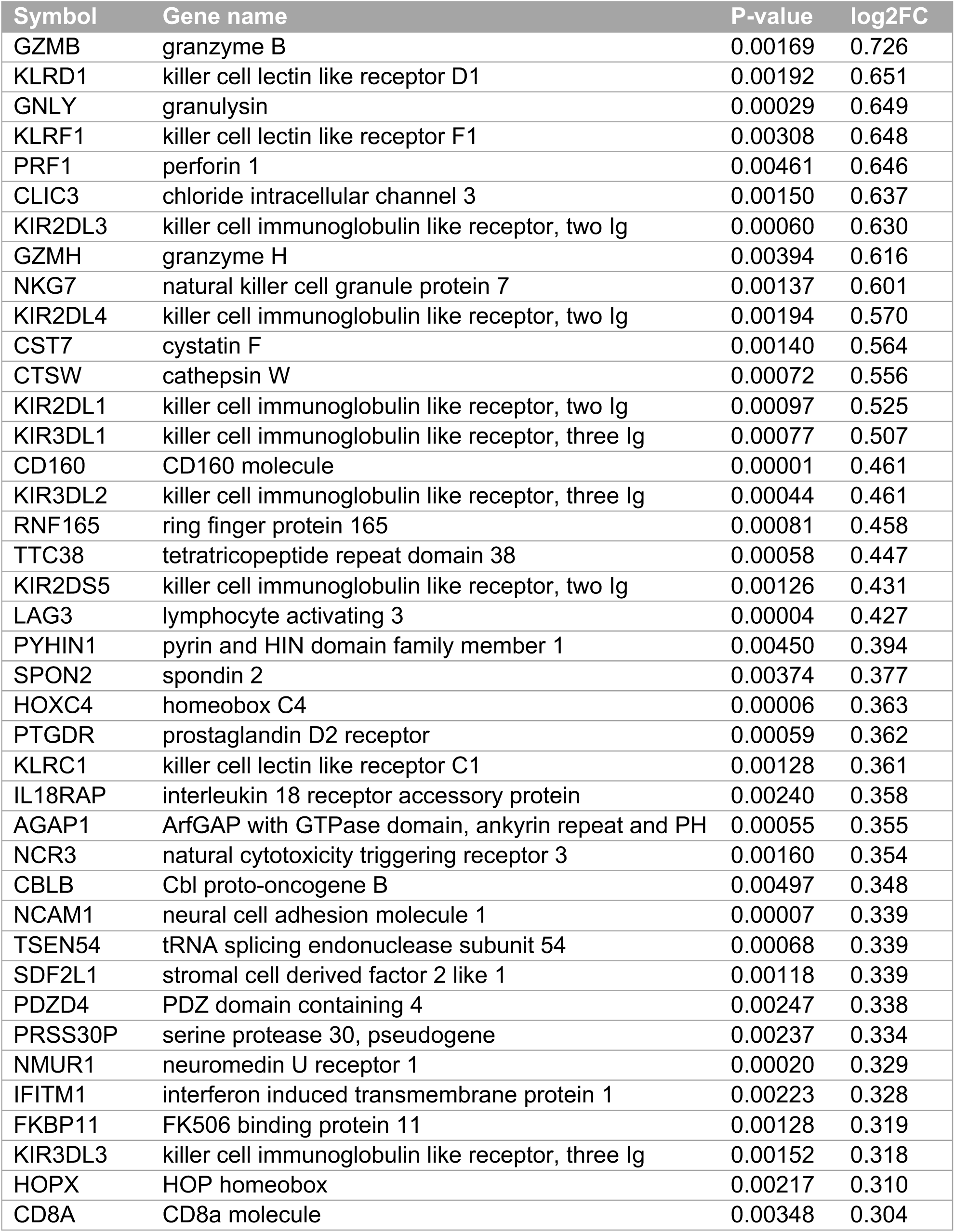
Top 40 of differentially expressed between post-treatment 3 months vs pre-treatment

| Symbol | Gene name | P-value | log2FC |
| --- | --- | --- | --- |
| GZMB | granzyme B | 0.00169 | 0.726 |
| KLRD1 | killer cell lectin like receptor D1 | 0.00192 | 0.651 |
| GNLY | granulysin | 0.00029 | 0.649 |
| KLRF1 | killer cell lectin like receptor F1 | 0.00308 | 0.648 |
| PRF1 | perforin 1 | 0.00461 | 0.646 |
| CLIC3 | chloride intracellular channel 3 | 0.00150 | 0.637 |
| KIR2DL3 | killer cell immunoglobulin like receptor, two Ig | 0.00060 | 0.630 |
| GZMH | granzyme H | 0.00394 | 0.616 |
| NKG7 | natural killer cell granule protein 7 | 0.00137 | 0.601 |
| KIR2DL4 | killer cell immunoglobulin like receptor, two Ig | 0.00194 | 0.570 |
| CST7 | cystatin F | 0.00140 | 0.564 |
| CTSW | cathepsin W | 0.00072 | 0.556 |
| KIR2DL1 | killer cell immunoglobulin like receptor, two Ig | 0.00097 | 0.525 |
| KIR3DL1 | killer cell immunoglobulin like receptor, three Ig | 0.00077 | 0.507 |
| CD160 | CD160 molecule | 0.00001 | 0.461 |
| KIR3DL2 | killer cell immunoglobulin like receptor, three Ig | 0.00044 | 0.461 |
| RNF165 | ring finger protein 165 | 0.00081 | 0.458 |
| TTC38 | tetratricopeptide repeat domain 38 | 0.00058 | 0.447 |
| KIR2DS5 | killer cell immunoglobulin like receptor, two Ig | 0.00126 | 0.431 |
| LAG3 | lymphocyte activating 3 | 0.00004 | 0.427 |
| PYHIN1 | pyrin and HIN domain family member 1 | 0.00450 | 0.394 |
| SPON2 | spondin 2 | 0.00374 | 0.377 |
| HOXC4 | homeobox C4 | 0.00006 | 0.363 |
| PTGDR | prostaglandin D2 receptor | 0.00059 | 0.362 |
| KLRC1 | killer cell lectin like receptor C1 | 0.00128 | 0.361 |
| IL18RAP | interleukin 18 receptor accessory protein | 0.00240 | 0.358 |
| AGAP1 | ArfGAP with GTPase domain, ankyrin repeat and PH | 0.00055 | 0.355 |
| NCR3 | natural cytotoxicity triggering receptor 3 | 0.00160 | 0.354 |
| CBLB | Cbl proto-oncogene B | 0.00497 | 0.348 |
| NCAM1 | neural cell adhesion molecule 1 | 0.00007 | 0.339 |
| TSEN54 | tRNA splicing endonuclease subunit 54 | 0.00068 | 0.339 |
| SDF2L1 | stromal cell derived factor 2 like 1 | 0.00118 | 0.339 |
| PDZD4 | PDZ domain containing 4 | 0.00247 | 0.338 |
| PRSS30P | serine protease 30, pseudogene | 0.00237 | 0.334 |
| NMUR1 | neuromedin U receptor 1 | 0.00020 | 0.329 |
| IFITM1 | interferon induced transmembrane protein 1 | 0.00223 | 0.328 |
| FKBP11 | FK506 binding protein 11 | 0.00128 | 0.319 |
| KIR3DL3 | killer cell immunoglobulin like receptor, three Ig | 0.00152 | 0.318 |
| HOPX | HOP homeobox | 0.00217 | 0.310 |
| CD8A | CD8a molecule | 0.00348 | 0.304 |

### Functional interpretation of transcriptomic changes induced by pazopanib

The top ten differentially modulated canonical pathways in post-treatment vs pre-treatment samples are shown in **Figure 2A-C.** Genes of the top 3 pathways in each comparison are shown in **Figure 2D**. The graphical representation of the top pathway at each time point comparison is shown in **Supplementary Figure 1**. The majority of the top canonical pathways modulated by pazopanib (7/10 in both *Post 3* and *Post 6* comparison) were associated with immune functions (**Figure 2A-D).** The perturbations induced at the 3^rd^ month of treatment are consistent with the triggering of NK/cytotoxic signaling, the positive modulation of the crosstalk between dendritic cells and NK cells, the regulation of IL-2, T cell receptor (TCR) signaling, and IL-8 signaling. After 3 further months of treatment, an attenuation of the immune modulatory effect induced by pazopanib was observed. This was substantiated by the downregulation of transcripts associated with T helper (Th)-1 and Th2 functional orientation when comparing *Post 6 vs Post 3* samples. The activation of NK-related pathways was still sustained at the 6^th^ montht of treatment, although attenuated.

**Figure 2:**
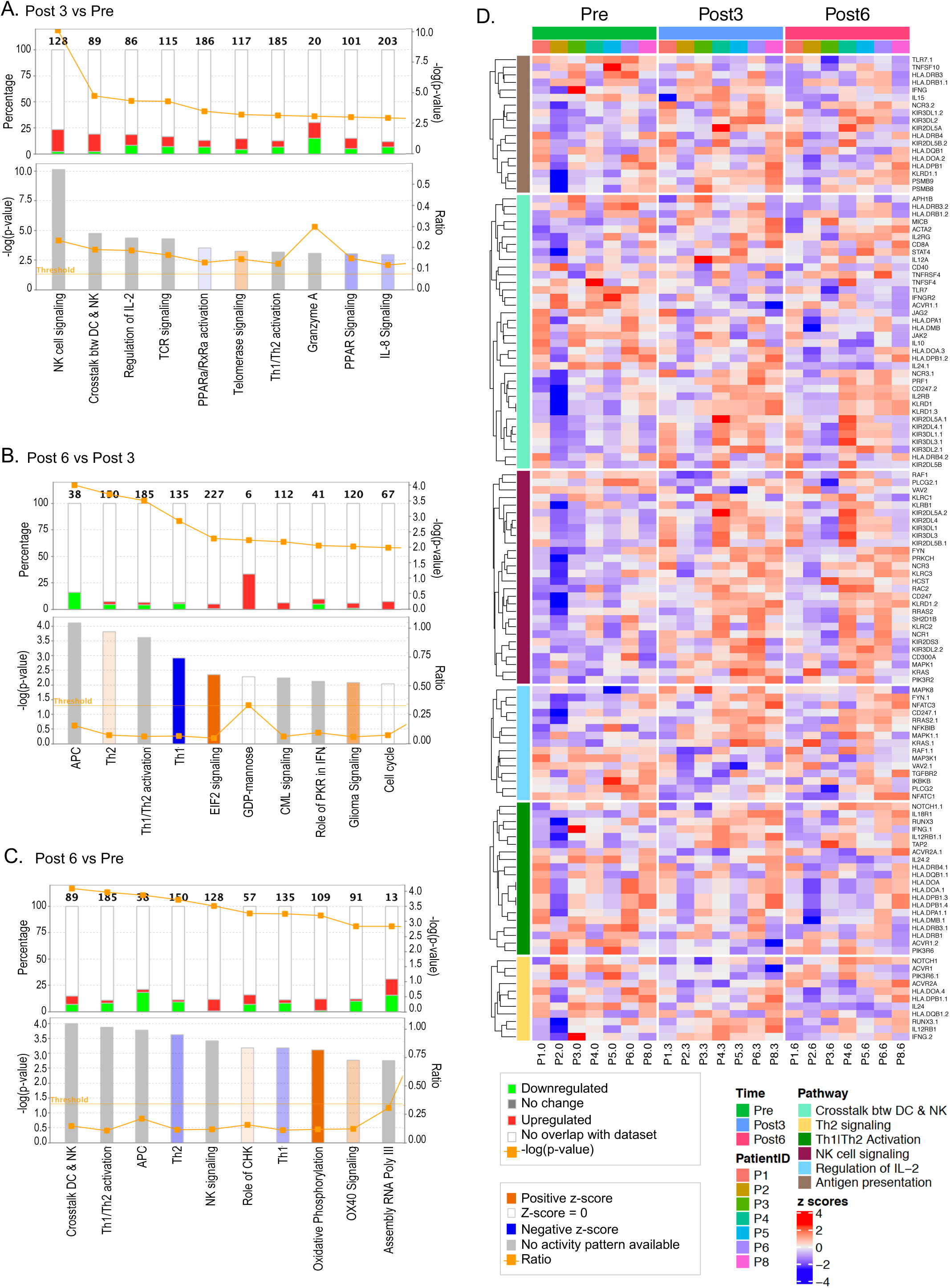
Impact of pazopanib treatment on gene expression. Top ten canonical pathways ranking modulated by treatment identified using IPA analysis according to significance level (*paired t-test, p* < *0.05*). (A) post-treatment 3 months (*Post 3*) vs baseline (Pre), (B) Post treatment 6 months (*Post 6*) vs *Post 3*, and (C) *Post 6* vs *Pre*. (D) Top 3 pathways genes (as listed in IPA software) in each timepoint comparison were presented by heatmap on the right panels. The z-score was used to indicate the direction of the pathway activation.

### Pazopanib-induced molecular fingerprints

We applied modular repertoire analysis to further dissect the immune-modulatory properties of pazopanib. The percentage of responsive transcripts constitutive of a given module was determined at each time point (see methods for details). The group comparison analysis confirmed that module perturbations peaked at 3 months of treatment and decreased at 6 months. These perturbations include upregulation of modules M3.6 and M8.46 defining cytotoxic/NK cells, M4.11 (plasma cells), and M8.89 (immune response). Moreover, the responsiveness of M4.14 (monocytes) was decreased (**Figure 3**). However, mapping perturbations of the modular repertoire for a group of subjects does not account for the heterogeneity observed at the individual level. We therefore performed deeper individual-level analysis. This approach demonstrated that pazopanib administration was associated with the decrease of M9.34 (immunosuppressive) in the majority of patients. The most coherent changes were represented by modulations of cytotoxic/NK cells modules (M3.6 and M8.46) while the majority of the other modules demonstrated a considerable heterogeneity. Interestingly, a rapid increase of Interferon (IFN) modules (M1.2 and M3.4) was observed exclusively in patients displaying up-regulation of cytotoxic/NK cells modules (**Figure 4, Supplementary Table 2)**

**Figure 3:**
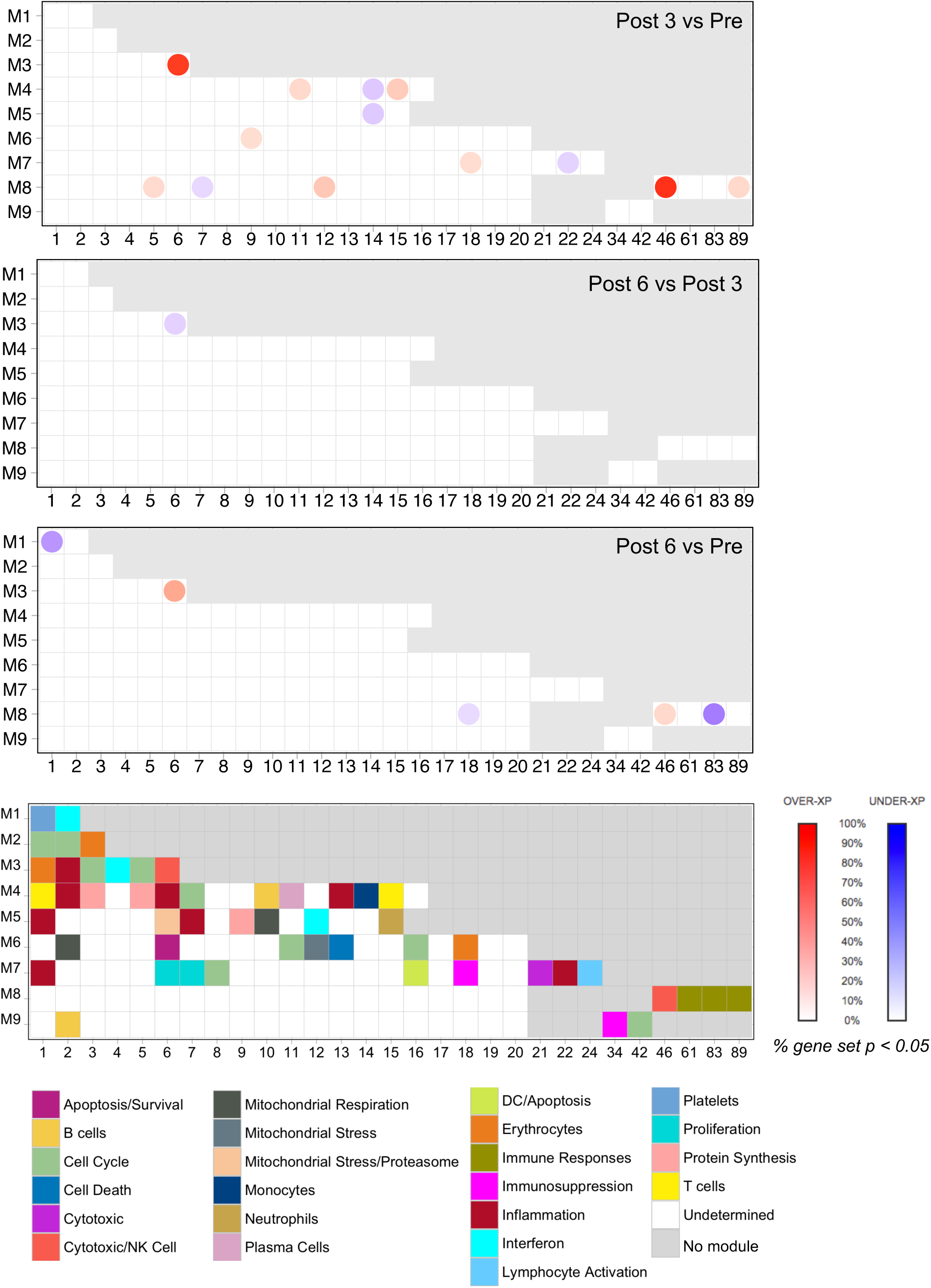
Modular mapping of changes in blood transcriptome elicited by pazopanib in mRCC patients. Changes in transcript abundance measured in PBMCs using whole-genome arrays were mapped against a pre-constructed repertoire of co-expressed gene sets (modules). The proportion of transcripts for which abundance was significantly changed in comparison between samples collected at 3 months (*Post 3*) vs baseline (*Pre*), 6 months (*Post 6*) vs 3 months (*Post 3*) and 6 months (*Post 6*) vs baseline (*Pre*) in each module. When the percentage of response exceeds 15 %, the module was considered as responsive to treatment. Responsive modules are mapped on a grid, the proportion of significant transcripts for each module is represented by a spot of color, with red representing increased abundance and blue representing decreased abundance. The degree of intensity of the spots denotes the percentage of transcripts in a given module showing significant difference in abundance in comparison to the baseline. A legend is provided with functional interpretations indicated at each position of the grid by a color code. Functional interpretations are indicated by the color-coded grid at the bottom of the figure.

**Figure 4:**
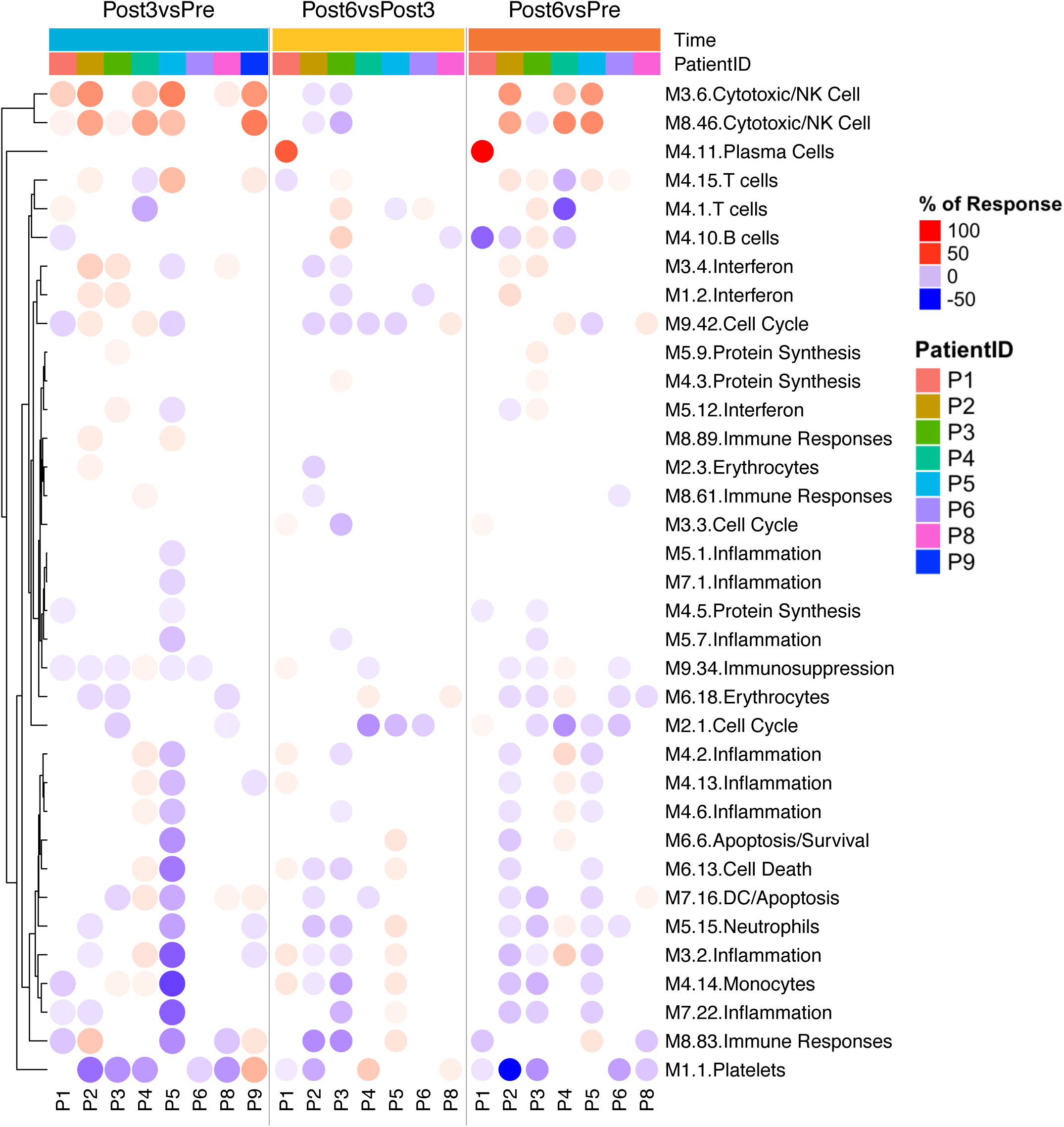
Mapping perturbations of the modular repertoire across individual samples. Individual comparisons. Percentage of response of Individual patients at *Post 3* vs *Pre* (blue), *Post 6* months vs *Post 3* months (yellow), and *Post 6* months vs *Pre* (orange). The expression profile for each individual patient was calculated as a FC and difference relative to an expression of individual samples in each time point. For determining post-treatment changes for individual subjects, a cut-off is set against which individual genes constitutive of a module are tested (|FC| > 1 and |difference| >10).

### Modulation of leukocyte functional orientation induced by pazopanib as derived by transcriptomic data

To estimate the changes in leukocyte populations, we used single sample Gene Set Enrichment Analysis (ssGSEA). Comparison of the enrichment scores of post-treatment and their baseline pre-treatment, showed that NKCD56^dim^, NKCD56^bright^, T gamma-delta (Tgd), NKT, cytotoxic cells and CD8 T cells increased coherently at 3 months of treatment and subsequently slightly decreased without reaching baseline levels (**Figure 5**). Conversely, regulatory T cells (Tregs) signature was decreased and a similar trend was observed for MDSCs (**Figure 5C**). These results suggest that pazopanib induces synergistic immune modulations by enhancing protective immunity and reducing suppressive mechanisms.

**Figure 5:**
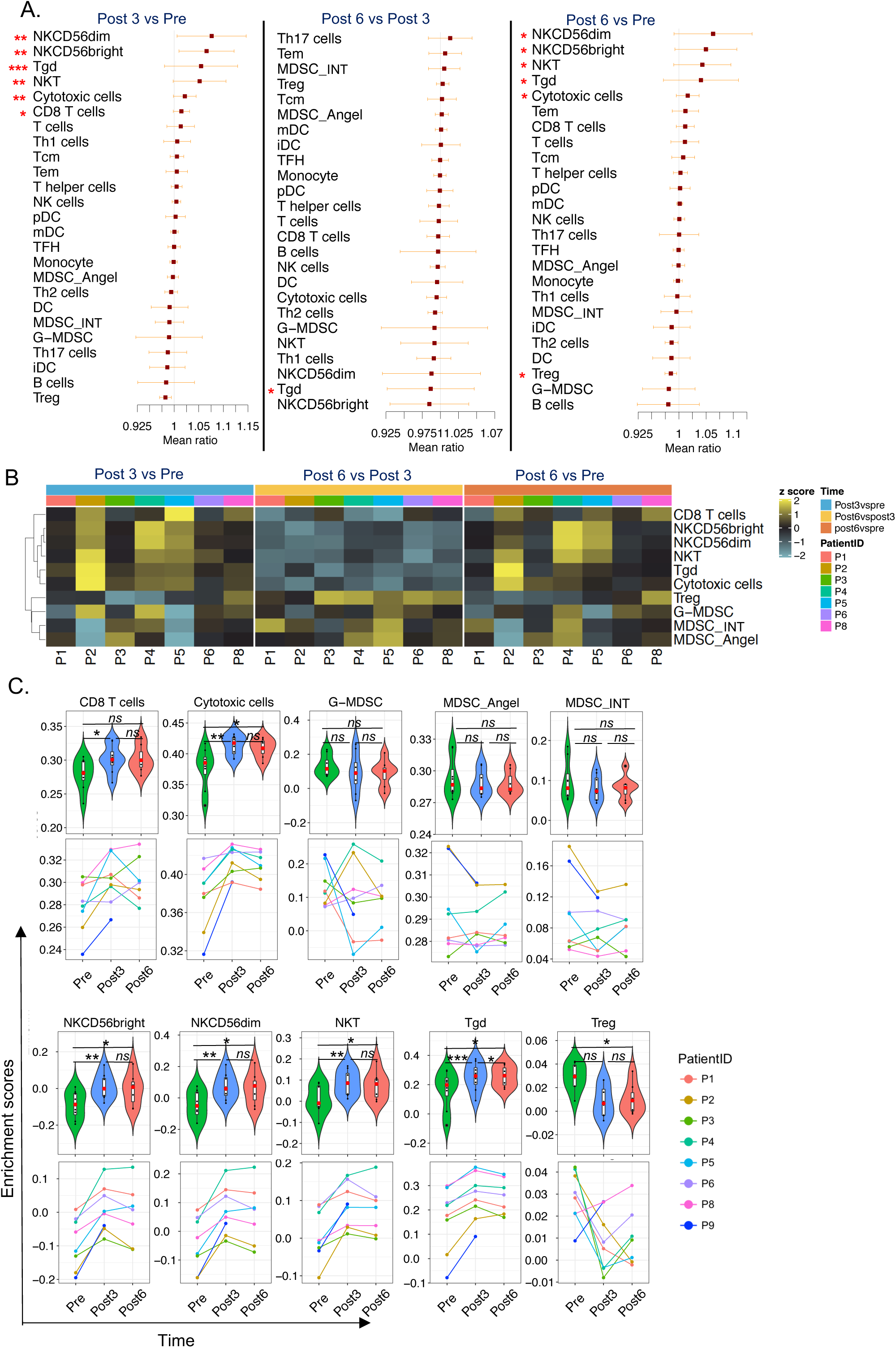
Cell-type specific analysis in pre-treatment and post-treatment samples. (A) Forest plot of leukocyte enrichment score comparison between *Post 3* vs *Pre, Post 6* vs *Post 3*, and *Post 6* vs *Pre*. (B) Heatmap analysis of fold-change of leukocyte enrichment score; the fold change-scored values of representative fold-change between *Post 3* vs *Pre, Post 6* vs *Post 3*, and *Post 6* vs *Pre* are displayed in a heatmap. (C) Violin plots and line charts of significant cell types. Red asterisks: * *= p* < 0.05, ** *= p* < 0.01, *** *= p* < 0.001; *ns = p* not significant.

### Flow cytometry analysis confirms the positive immune modulation associated with pazopanib administration

Transcriptome profiling in bulk cell populations provided a high-level and unbiased perspective on the changes taking place following initiation of therapy. It is ideally complemented by flow cytometry analyses which provide a targeted but highly granular view of changes taking place at the cellular and protein levels.

Multicolor flow cytometry analysis of PBMCs was performed concomitantly in biological replicates of the same blood samples submitted to transcriptional profiling, plus an additional patient for whom RNA was not available. A broad panel of markers encompassing innate and adaptive immune cell subsets of the lymphoid and myeloid repertoire was studied and modulation in on-treatment with respect to baseline samples was assessed. The analysis showed that pazopanib administration was associated with a remarkable increase of activated and cytolytic effectors including the subset of activated T cells (CD3^+^PD-1^+^), reported to contain tumor-specific T cells [40], activated NK cells (CD3^-^CD16^+^CD56^+^PD-1^+^) and cytotoxic NK cells (CD3^-^CD16^+^CD56^dim^) (**Figure 6**) [41]. Of note, this evidence is in line with the data that emerged from the transcriptional profiling, depicting an overall boost of genes involved in TCR signaling, cytotoxic cell populations and NK activity. Again, similarly to findings obtained via transcriptomic analyses, the detected changes over baseline were more evident at 3 months of therapy and tended to a plateau or decreased at 6 months. The boost of T and NK cell activation was paralleled by a significant decrease in the frequency of different myeloid cell subsets, including CD14^+^ monocytes, MONO-MDSCs (CD14^+^HLA-DR^neg^) [42], inflammatory monocytes (CD14^+^PDL-1^+^) [43], and PMN-MDSCs (CD15^+^) [42](**Figure 6A).** Regulatory T cells (CD4^+^CD25^high^Foxp3^+^) also displayed a remarkable reduction during treatment. Down-modulation of all these cell subsets was mostly detectable at 3 months during treatment with respect to baseline, with a stabilization of monocytic MDSCs or/and a rebound for total and inflammatory monocytes in cell frequencies at 6 months (**Figure 6**). Taken together, the kinetics of immune modulation as detected by flow cytometry are in line with the data emerging from the transcriptional profiling data and confirm the transient nature of immunomodulation mediated by pazopanib.

**Figure 6:**
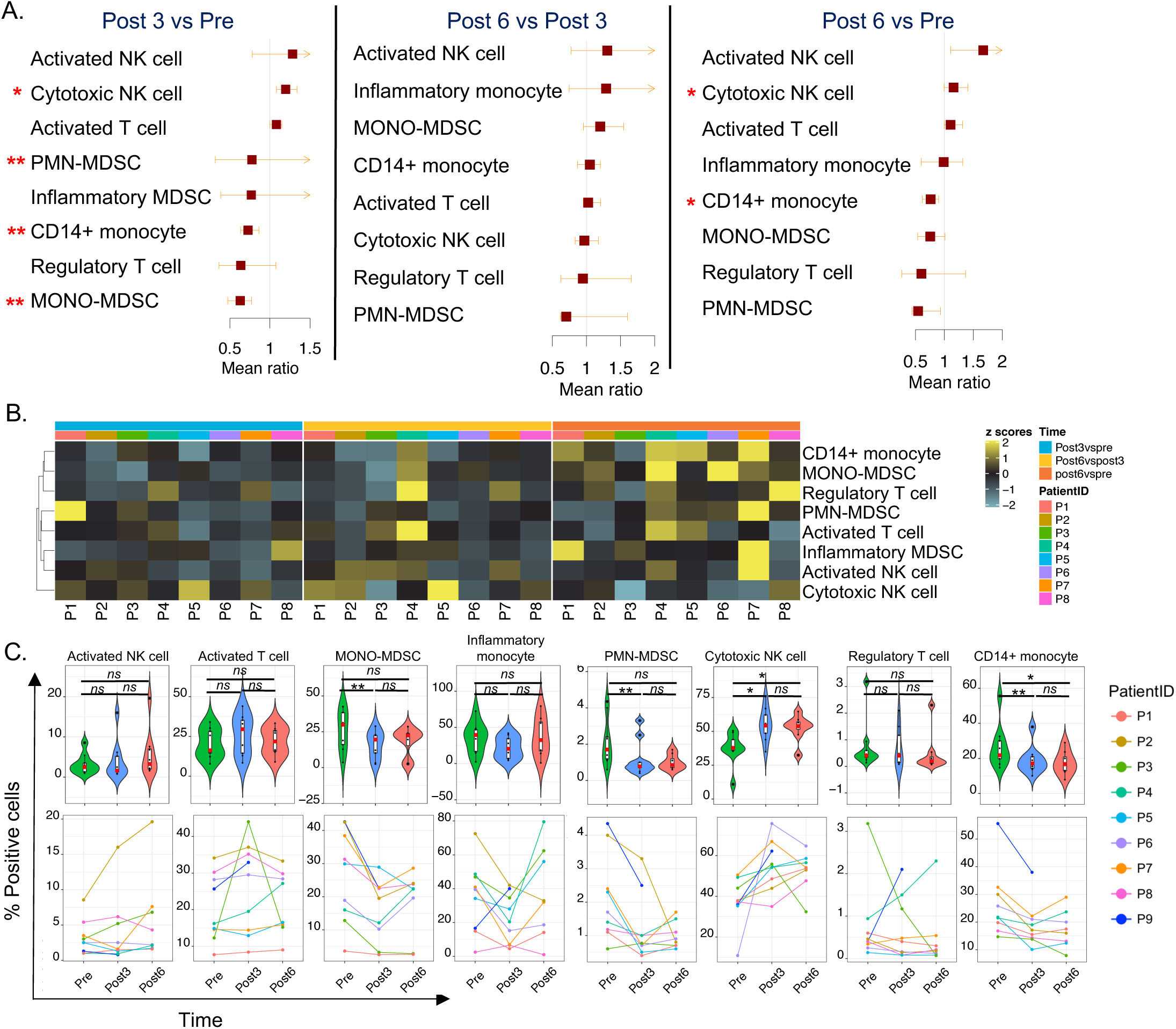
Flow cytometry analysis in samples with pre-treatment and post-treatment. (A) Forest plot of the ratio of cell-type proportions between *Post 3* vs *Pre, Post 6* vs *Post 3*, and *Post 6* vs *Pre* analyzed by flow cytometry. (B) Heatmap analysis of fold-change of cell-type proportions; the z-scored values of representative fold-change between *Post 3* vs *Pre, Post 6* vs *Post 3*, and *Post 6* vs *Pre* are displayed in a heatmap. (C) Violin plots and line chart of significant cell types. Red asterisks: * *= p* < 0.05, ** *= p* < 0.01, *** *= p* < 0.001; *ns = p* not significant.

### Intratumoral estimates of MDSC are associated with poor prognosis in kidney cancer

One of the more remarkable findings obtained through combined transcriptomic and flow-cytometry based immune monitoring is the decrease in MDSC populations and associated signatures.

To explore the relevance of our observation, and as no data exists regarding the prognostic role of MDSCs in kidney cancer, we assessed the expression of the three MDSC signatures in The Cancer Genome Atlas (TCGA) clear cell RCC cohort (KIRC, N=517, **Figure 7)**. The MDSC_INT signatures was strongly associated with decreased overall survival (MDSC_INT High vs LowMed, HR =2.057 (95% CI = 1.52-2.79, **Figure 7A)**. In particular, MDSC High group had poor prognosis, while MDSC Low and Med group (**Supplementary Figure 2**) have similar favorable prognosis. No such differences were observed using the other two MDSC signatures MDSC_Angel and G-MDSC, suggesting that MDSC_INT, which was developed experimentally based on extracellular-driven monocyte-MDSC differentiation, might represent a novel prognostic biomarker in kidney cancer. Remarkably, MONO-MDSCs were strongly suppressed after pazopanib treatment **(Figure 6).** MDSC_INT correlates with Stage and Grade, which are major prognostic factors in kidney cancer **(Figure 7B)**. We then assessed the relationship between MDSC_INT with the disposition of oncogenic pathways, and found that MDSC_INT expression linearly correlates with many oncogenic processes associated with cancer aggressiveness, including angiogenesis (R = 0.59, *p* < 2e-16), and epithelial-to-mesenchymal transition (EMT) (R = 0.75, *p* < 2e-16, **Figure 7C-D)** although there was no overlapping between MDSC_INT signature and angiogenesis or EMT (**Supplementary Figure 3**). Despite the correlation with Stage and Grade, MDSC_INT retained its prognostic value even when included in a Cox regression multivariable model (**Figure 7E**).

**Figure 7:**
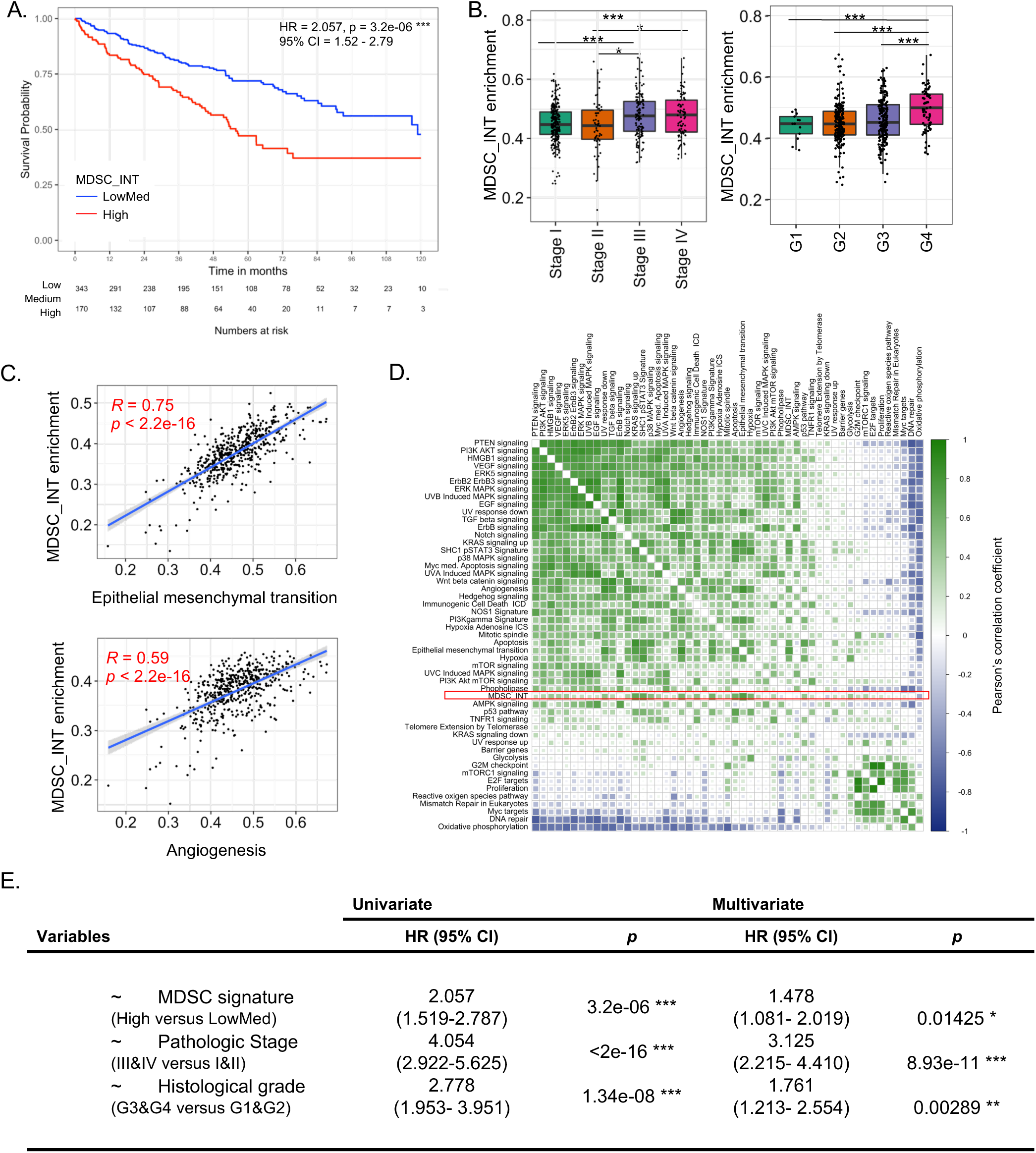
Prognostic implications of MDSC gene signature in kidney renal clear cell carcinoma (KIRC, n = 515). (A) Kaplan Meier curves showing overall survival (OS) of patients within the highest tertile of MDSC_INT enrichment versus the two lower tertiles (LowMed). Cox proportional hazards statistic are shown. (B) Boxplots of MDSC_INT enrichment scores by AJCC pathologic stage (left) and histological grade (right). T-test: * *= p* < *0.05*, ** *= p* < *0.01*, *** *= p* < *0.001*. C) Scatterplots showing the association between MDSC_INT scores and the enrichment score (ES) of genes related to epithelial mesenchymal transition (upper), and angiogenesis (lower). Regression line with corresponding Pearson’s correlation coefficient (R) and p-value are shown. (D) Pearson correlation matrix of enrichment scores of tumor-associated pathways. MDSC_INT signature is indicated with a red square. (E) Univariate and multivariate overall survival Cox proportional hazards regression analysis including MDSC_INT signature, pathologic stage (III&IV versus I&II), and histological grade (G3&G4 versus G1&G2).

Thus, the analysis of the tumor specimens from the TCGA cohort permitted to expand our initial finding by providing indirect evidence about the role of MDSCs in mRCC progression. Indeed, this in turn suggests that MDSC suppression by pazopanib may be one of the means by which treatment could contribute to improved outcome.

## Discussion

This is the first translational study that investigates longitudinally the immunomodulatory effects of an anti-angiogenic therapy on PBMCs of mRCC patients through transcriptomic analysis. Our results show that pazopanib triggers cytotoxic cells and IFN pathways and relieves immunosuppression by reducing MDSCs. This invigoration of anti-tumor immunity was mostly evident after 3 months of therapy and still detectable, although less accentuated, at the 6^th^ month of treatment. The transcriptional profiling of PBMCs clearly revealed treatment-induced immunomodulation, detecting modifications validated by flow cytometry, but also expanding them by revealing pathway networks and broader functional information. Our data indicates that the analysis of transcriptional profiles of blood cells with the support of appropriate deconvolution approaches represents a valid strategy for monitoring immune cell behavior at a high throughput and reliable level.

To achieve the results reported here, one of the major challenges was the identification of gene-signatures appropriate to capture the activity of circulating MDSCs. Indeed, monocytic MDSCs are defined in flow cytometry only by the lack/low expression of HLA-DR in cells expressing the monocytic marker CD14, alone or in combination with CD11b and CD33 [42], but their genomic features have been poorly defined. To this aim, we used the dataset selected by Angelova [44] and Fridlender [45] as reference transcriptional data for myeloid cells and the data set obtained from human MDSCs generated *in vitro* according to a model developed in our laboratory [46]. This MDSC model was produced by exposing blood CD14^+^ monocytes to tumor extracellular vesicles, a process leading to cells highly overlapping for phenotype, immunosuppressive function and transcriptional profiles with MDSCs isolated from blood of melanoma patients [46–48]. The gene signature reflects most of the signaling pathways expected for these cells and overlap with monocytes sorted from cancer patients [46]. Our MDSC signature, applied to bulk tumors, was the only one with prognostic implications in mRCC, confirmed in multivariate analyses, providing here an essential contribution for the estimations of MDSCs in different tissues.

Pazopanib is a multikinase inhibitor, among the ones of standard of care for first-line treatment of mRCC patients, especially if they are “low risk” according to Heng criteria. This TKI targets the VEGF receptor, platelet-derived factor receptor (PDGFR), KIT (proto-oncogene receptor tyrosine kinase), fibroblast growth factor receptors (FGFR) and RAF kinases [36–38]. The broad array of targets largely justifies the multifaceted immunomodulating activity we observed in our study, which involved different myeloid cell subsets together with T cells, NK cells and Tregs. However, given the known inhibitory activity of several antiangiogenics on lymphocytes when tested *in vitro* [49], it is tempting to speculate that this general immunological reshaping mostly stems from blunting of the complex immunosuppressive pathways mediated by MDSC and Tregs. Indeed, pazopanib might exert off-target inhibitory effects on MDSCs by acting on the VEGFR down-stream signaling [50, 51], or by interfering with the KIT pathway, another key node in myeloid cell function favoring DC reprogramming [8]. Hence, the boost of T and NK cell activation and cytolytic functions we broadly observed by both transcriptomics and flow cytometry, likely results from reduced intratumoral immunosuppression leading to an enhanced anti-tumor immune response, rather than from a stimulatory activity of the drug on these immune cell subsets.

A recent study demonstrated that pazopanib induces DC maturation *in vitro* [52], while novel VEGF-directed drugs, such as axitinib, enhances the expression of NKG2D ligand and consequently potentiates NK-cell cytolytic activity [53].

Previous studies in humans have shown that the anti-VEGF monoclonal antibody sunitinib reduces the abundance of monocytes [54] (assessed 4 and 6 weeks post-treatment), monocytic MDSCs [55] and Tregs [56] (assessed at 4 weeks after treatment only) in the blood. However, the persistence of Tregs suppression (assessed at 1.5 and 3 months post-treatment) has been correlated with prolonged overall survival [57]. There are no studies that have assessed immunologic perturbations in patients receiving pazopanib as first line treatment. Only one study has assessed immunologic changes in mRCC patients treated with pazopanib, but administered as third-line treatment, therefore studying patients with a potentially heavily compromised immune system. In such work, Pal *et al* [58] observed that post-treatment non-responder patients had lower levels of HGF, VEGF, IL-6, IL-8 and soluble IL-2R, and increased numbers of monocytic MDSCs as compared with responders. Failure to detect an overall decrease of MDSCs as compared to baseline might result from the fact that analyses were performed at late time points (at 6 and 12 months after treatment initiation). In fact, our study showed that the changes induced by pazopanib partially revert after 6 months.

The number of patients analyzed here was rather small but reflected the rarity of RCC patients who could be prospectively enrolled for first-line pazopanib administration especially in research hospitals with competitive clinical trials enrolling. Even so, dynamic changes were extremely coherent across patients and confirmed using orthogonal immune monitoring platforms and analyses. The data reported here provide a set of key information that might have relevant implications for the design of combinatory treatment strategies in mRCC clinical setting. First, pazopanib mediates a specific reshaping of tumor immunity that should favor a prompter response to immunotherapy due to the decrease of immunosuppressive effectors and the concomitant boost of PD-1^+^ T cells and NK cells. Secondly, this effect reaches its peak at the 3^rd^ month of treatment but tends to be attenuated at later time points likely due to the homeostatic mechanisms that regulate systemic immunity and tumor-mediated immunosuppression. These data indicate that a short-term pre-conditioning treatment with pazopanib might create the optimal immune setting for ICB to potentiate antitumor immune responses *in vivo*. In this view, the combination of pazopanib with PD-1 blockers, which resulted so far in unmanageable toxicity during initial clinical testing [59], could be replaced by intermittent schedules that might help maximizing treatment synergies. The concomitant administration of PD-1/PD-L1 inhibitors with anti-angiogenic monoclonal antibodies (bevacizumab, IMmotion 151 trial) [60] or TKI axitinib (MK426 [61] and JAVELIN RENAL101 [62] trials), is rapidly emerging as a strategy to increase overall survival in RCC patients, with further ongoing studies evaluating combination with novel TKI lenvatinib (CLEAR trial), cabozantinib (CheckMate 9ER), and tivozanib [63] but the optimal schedule of such combinations still needs to be defined. In this context, our data suggest that more dynamic and innovative approaches based on intermittent or alternate schedules could be also explored to ameliorate the therapeutic index of combinatorial regimens in cancer [64, 65].

## Conclusion

Transcriptional profiling of blood immune cells, particularly if combined with specific deconvolution programs to finely dissect the behavior of diverse immunological components, represents a reliable tool to readily capture the immunomodulating properties of anticancer therapies for designing scientifically-sound combination therapies. Thanks to this analysis we found that pazopanib has a strong immune modulatory effect peaking at the 3^rd^ month of treatment and consists in relieved immunosuppression with enhanced cytotoxic T and NK mechanisms and IFN pathways. Taken together with the detrimental role of MDSCs in mRCC demonstrated in the TCGA cohort, these results provide a strong rationale for the use of TKIs as preconditioning strategy to improve immunological and clinical efficacy of immunotherapy in cancer patients.

## Materials and Methods

### Patients and study description

From January 2016 to June 2016 nine patients (8 males, 1 female) with metastatic clear cell RCC were treated with first line pazopanib as per clinical practice in Istituto Nazionale dei Tumori, Milan, Italy. Safety assessment included physical examination and laboratory tests every month. All patients had a good performance status (ECOG 0:8/9, ECOG 1: 1/9), a median age of 65 years and prevalence of intermediate risk according to Heng score (5/9). They received pazopanib at a standard dose of 800 mg orally once daily, continuously, for at least 6 months. All patients signed an informed consent according to a protocol approved by the INT Ethical Committee [INT146/14].

### Blood collection

Blood samples (30 ml) were obtained from 9 patients at baseline (*Pre*), and at the 3^rd^ and 6^th^ month during therapy (*Post 3* and *Post 6*). For one single patient, samples were collected only at baseline and at 3 months therapy. Blood was processed within 1 hour from withdrawal. PBMCs were separated by Ficoll gradient (Leuco-sep tubes, ThermoFisher Scientific) and viable cells stored in liquid nitrogen until use, or frozen in Qiazole (Qiagen) for RNA extraction and gene expression profiling.

### Transcriptomic analysis

Suitable material for transcriptional analysis was available from 8 patients. RNA was extracted using miRNAeasy kit (Qiagen). After quality check and quantification by 2100 Bioanalyzer system (Agilent) and Nanodrop ND-1000 spectrophotometer (ThermoFisher), respectively, RNA expression was assessed using Illumina HT12v4 BeadChip. Illumina’s BeadStudio version 1.9.0 software was used to generate signal intensity values from the scans. Data were further processed using the Bioconductor “Lumi” package. Following background correction and quantile normalization, expressions were log2-transformed for further analysis. Raw expression and normalized data matrix have been deposited at NCBI’s Gene Expression Omnibus database (http://www.ncbi.nlm.nih.gov/geo/), with accession numbers GSE146163. From a total of 47,323 probes arrayed on the Illumina HT12v4 beadchip, the probes targeting multiple genes were collapsed (average expression intensity) and a final data matrix containing 12,913 unique genes was generated. Data analyses were performed using R (Version 1.0.44, RStudio Inc.) and Ingenuity Pathway Analysis (QIAGEN Bioinformatics). The comparison between each group (*Post 3 vs pre, Post 3 vs Post 6* and *Post 6 vs Pre*) was performed using the paired *t*-test. For detection of differentially expressed genes we used a *p* value cutoff of 5×10^−3^, and false discovery rate (FDR) was provided as descriptive statistic (**Supplementary Table 1**), and not to dictate significance as the risk of type I error was mitigated by the use of orthogonal platforms (i.e., flow cytometry) for validation purposes. Hierarchical clustering was performed using the function “Heatmap” from the R package “ComplexHeatmap” [66]. Euclidean distance and complete linkage methods were used by default. Principle component analysis (PCA) was performed using the R function “scatterplot3d” package. The first three principal components, PC1, PC2 and PC3, were plotted against each other.

#### Pathway analysis

Gene ontology analyses were performed using Ingenuity Pathway Analysis (QIAGEN Bioinformatics). A relaxed *p* value cut-off of 0.05 was used to select transcripts for pathway analysis. The proportion of upregulated and downregulated transcripts was represented. The z-score was used to indicate the direction of pathway deregulation. Transcripts from the top three pathways in each comparison group were plotted in the corresponding heatmaps.

#### Leucocyte subset estimations

To estimate the enrichment of various cell types, gene expression deconvolution analyses were performed with ssGSEA [1, 2] implemented in the “GSVA package” using cell-specific signatures (**Supplementary Table 3**): T cells, CD8 T cells, cytotoxic T cells, T helper 1 cells (Th1 cells), central memory T cells (Tcm), effector memory T cells (Tem), T helper cells, T follicular helper cells (Tfh), Th2 cells, Th17 cells, gamma delta T cells (Tgd), natural killer cells (NK cells), NK CD56^dim^, NK CD56^bright^ cells, B cells, neutrophils, eosinophils, mast cells [67], regulatory T cells (Treg), NKT cells, monocytes, dendritic cells (DC), immature DCs (iDC), plasmacytoid DCs (pDC), myeloid DCs (mDC) [44]. For MDSCs, we constructed a specific signature based on 25 genes highly correlated in the present dataset, selected from the top 100 genes upregulated in extracellular vesicle (EV)-MDSCs vs monocytes (MDSC_INT, **Supplementary Figure 4**) in our recent work [46]. Additional MDSC signatures include the one proposed by Angelova *et al.* [44] (MDSC_Angel), based on markers selected according to the literature, and a granulocytic myeloid-derived suppressor cell (G-MDSC) signature defined by comparing G-MDSC vs naïve neutrophils [45]. Forest plots were plotted by using mean enrichment scores (ESs) ratio between *Post 3 vs Pre, Post 6 vs Post 3* and *Post 6 vs Pre*. Differentially expressed ESs between pre-treatment and post-treatment were calculated through paired *t-*test (*p* < 0.05).

#### Modular repertoire analysis

A set of 260 modules (co-expressed genes) was used for the analysis of this data set. This fixed modular repertoire was a priori determined, being constructed based on co-expression measured across 9 reference datasets encompassing a wide range of diseases (infectious, autoimmune, inflammatory) [17, 18, 68] (https://github.com/Drinchai/DC_Module_Generation2). This data-driven approach allowed the capture of a broad repertoire of immune perturbations which were subsequently subjected to functional interpretation. This collection of annotated modules was then used as a framework for analysis and interpretation of our blood transcriptome dataset. The approach used for the construction, annotation and reuse of modular blood transcriptome repertoires was previously reported [14, 15, 17, 18, 68]. After normalization, raw expression intensity was used for the module analysis. Briefly, data was transformed from gene level data into module (M) level activity scores, both for group comparison *(Post 3 vs Pre, Post 6 vs Post 3* and *Post 6 vs Pre)* and individual patients comparison at each time point. The modules defined by this approach (M1–M9, a total of 260 modules) were used as a framework to analyze and interpret this dataset. For group comparisons, the expression profile at each time point was calculated as a FC relative to a mean expression of all samples within that time points. Then, paired *t-* test was used to evaluate each time point comparison. If the FC between each group comparison was greater than 1, and the *p* value < 0.05, the transcript was considered as upregulated. If the FC between each group comparison was less than 1, and the *p* value < 0.05, it was considered as downregulated. Then the percentages of “module responsiveness” were calculated for each module. For individual comparison, the expression profile for each individual patient was calculated as a FC and difference relative to the expression of individual samples at each time point. If the FC between each time point comparison was more than 1, and difference more than 10, the transcript was considered as upregulated. If the FC between each time point comparison was less than 1, and the difference less than 10, it was considered as downregulated. For both, group and individual comparisons, the “module-level” data is subsequently expressed as a percent value representing the proportion of differentially regulated transcripts for a given module. A module was considered to be responsive when more than 15% of the transcripts were down or upregulated.

### Multiparameter flow cytometry

PBMC samples from 9 patients were thawed and tested simultaneously for all time points by flow cytometry. Phenotypic profiling was performed after labeling PBMCs with monoclonal fluorochrome-conjugated antibodies: CD14 FITC (Clone M5E2, BD Pharmingen), CD3 FITC (Clone UCHT1, BD Biosciences) or KO525 (Clone UCHT1, Beckman Coulter), PD-1 APC (Clone MIH4, BD Pharmingen) or PC7 (Clone PD1.3, Beckman Coulter), HLA-DR APC (Clone G46-6, BD Pharmingen), CD15 PerCP-CY5.5 (Clone HI98, BD Pharmingen), PD-L1 PE (Clone MIH1, BD Pharmingen), CD4 PE (Clone RPA-T4, BD Pharmingen), CD25 PerCP-Cy5.5 (Clone M-A251, BD Pharmingen), CD56 ALEXA750 (Clone N901, Beckman Coulter), CD16 BV650 (Clone 3G8, BD Biosciences), Live/Dead Fixable Violet (ThermoFisher), FOXP3 APC (Clone FJK-16s eBioscience) used after cell permeabilization with the kit Perm Buffer (10X) and Fix/Perm Buffer (4X) (BioLegend), according to manufacturer’s instructions. Samples were incubated with Fc blocking reagent (Miltenyi Biotec) for 10 minutes at room temperature before the addition of monoclonal antibodies for 30 minutes at 4°C. Thereafter, samples were washed, fixed acquired by Gallios Beckman Coulter FC 500 or BD FACSCalibur (BD Biosciences) flow cytometers, and analyzed with Kaluza software (Beckman Coulter). Gating strategies are depicted in **Supplementary Figure 5**. Distinct cell subsets were quantified in terms of frequency rather than absolute numbers, since the latter are influenced by sampling manipulation procedures that are unrelated to biological patterns. Pre-treatment and post-treatment samples (*Post 3* vs *Pre, Post 6* vs *Post 3* and *Post 6* vs *Pre*) were compared by using paired *t-* test.

### TCGA transcriptomic Analysis

RNA-seq data from TCGA clear cell RCC (KIRC) cohort were downloaded using TCGA Assembler (v2.0.3). Data normalization was performed within lanes, and between lanes using R package EDASeq (v2.12.0) and quantile normalized using preprocessCore (v1.36.0). A single primary tumor sample was included per patient using the TCGA Assembler “ExtractTissueSpecificSamples” function. Previously flagged samples that did not pass assay-specific QCs were excluded [69]. Data was log2 transformed with an (+1) offset. Enrichment scores (ES) were calculated by ssGSEA on the log2 transformed, normalized gene-level data. Gene sets to define ES of tumor-associated pathways (n=51) were used as described previously [70]. The correlation between tumor-associated signatures was calculated using Pearson test and plotted using “corrplot” (v0.84).

### Survival Analysis

Clinical data from the TCGA-KIRC cohort was obtained from the TCGA Pan-Cancer Clinical Data Resource [71]. Patients were divided in tertiles based on enrichment scores of MDSC gene signatures (MDSC_INT, G-MDSC, and MDSC_Angel). Overall survival (OS) was used to generate Kaplan-Meier curves using a modified version of the ggkm function [72]. Survival data were censored after a follow-up period of 10 years. Hazards ratios (HR) between groups, corresponding *p* values, and confidence intervals were calculated using cox proportional hazard regression with R package survival (v2.41-3).

## Abbreviation List

DC: Dendritic cell
iDC: Immature dendritic cell
mDC: Myeloid dendritic cells
pDC: Plasmacytoid dendritic cells
ICB: Immune checkpoint blockade
IFN: Interferon
IPA: Ingenuity Pathway Analysis
MDSC: Myeloid derived suppressor cells
G-MDSC: Granulocytic Myeloid derived suppressor cells
M-MDSC: Monocytic Myeloid derived suppressor cells
mRCC: Metastatic renal cell carcinoma
NK: Natural killer cell
PCA: Principal component analysis
PBMCs: Peripheral blood mononuclear cells
Tcm: Central memory T cell
Tem: Effector memory T cells
Tfh: T follicular helper cells
Th2 cells: T helper 2 cells
Th1 cells: T helper 1 cells
Th17 cells: T helper 17 cells
Tgd: T gamma delta cells
Treg: Regulatory T cell
TKI: Tyrosine-kinase inhibitor
VEGF: Vascular endothelial growth factor

## Acknowledgements

This work was supported by the Associazione Italiana per la Ricerca sul Cancro (AIRC) Special Program Innovative Tools for Cancer Risk Assessment and early Diagnosis 5X1000 (no. 12162 to LR) and by Fondi 5×1000 Ministero della Salute 2015 (D/17/1VH to VH) and by the Italian Ministry of Health (RF-2016-02363001 to G.P.) and by Horizon 2020 PRECIOUS Project, Grant Agreement n. 686089 to LR. We would like to thank the patients and participating study team for making this study happen. DB, DR, and DC work has been supported by Sidra Medicine, Member of Qatar Foundation.

---

**Supplementary Figure 1:**
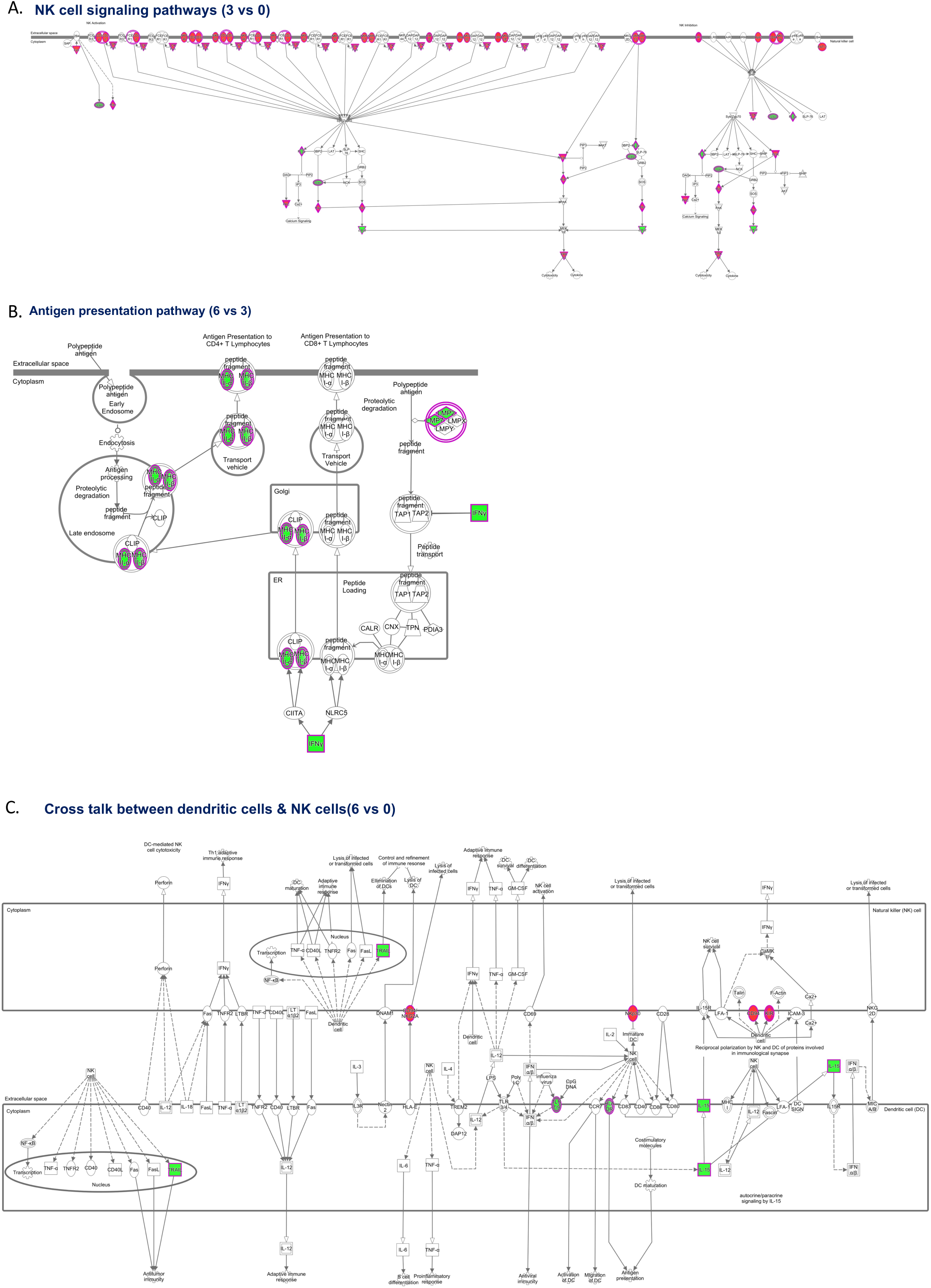
Top pathway in each time point comparison.

**Supplementary Figure 2:**
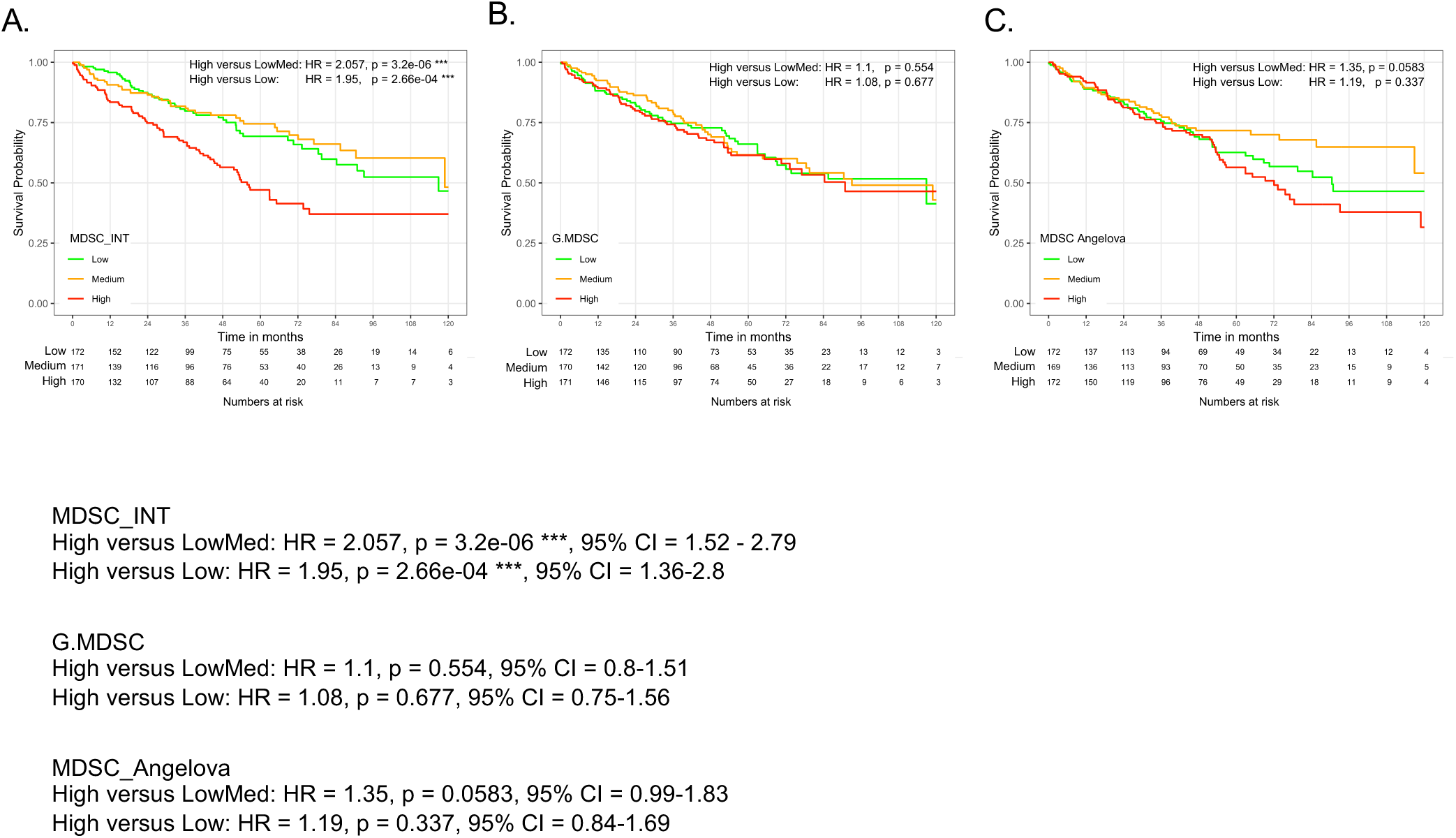
Prognostic value of MDSC signatures. Kaplan Meier curves showing overall survival (OS) of patients within tertiles of MDSC_INT (A), granulocytic MDSC (B), and MDSC Angelova (C) enrichment scores. Cox proportional hazards statistics of the high versus LowMed (combination of intermediate and low tertiles) and High versus Low tertiles are shown.

**Supplementary Figure 3:**
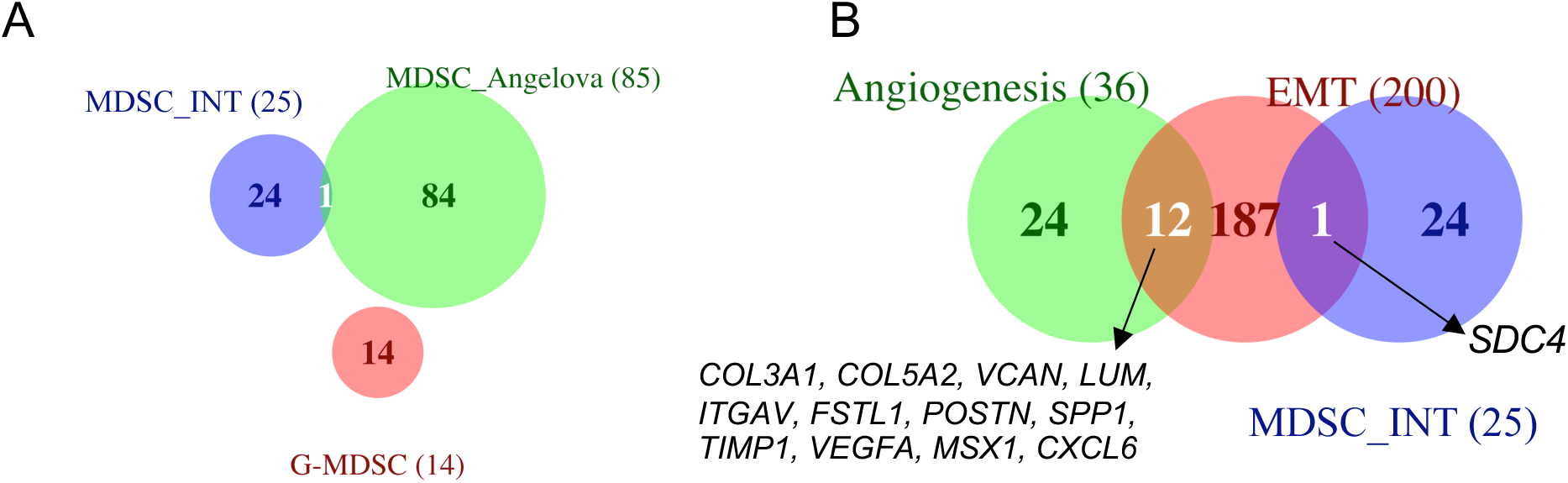
Venn diagrams of intersection between MDSC signatures; MDSC_INT, MDSC_Angelova and G-MDSC (A), and MDSC_INT and Cancer hallmark pathways (Angiogenesis and Epithelial mesenchymal transition) (B). The total number of genes in each area is reported between the parentheses.

**Supplementary Figure 4:**
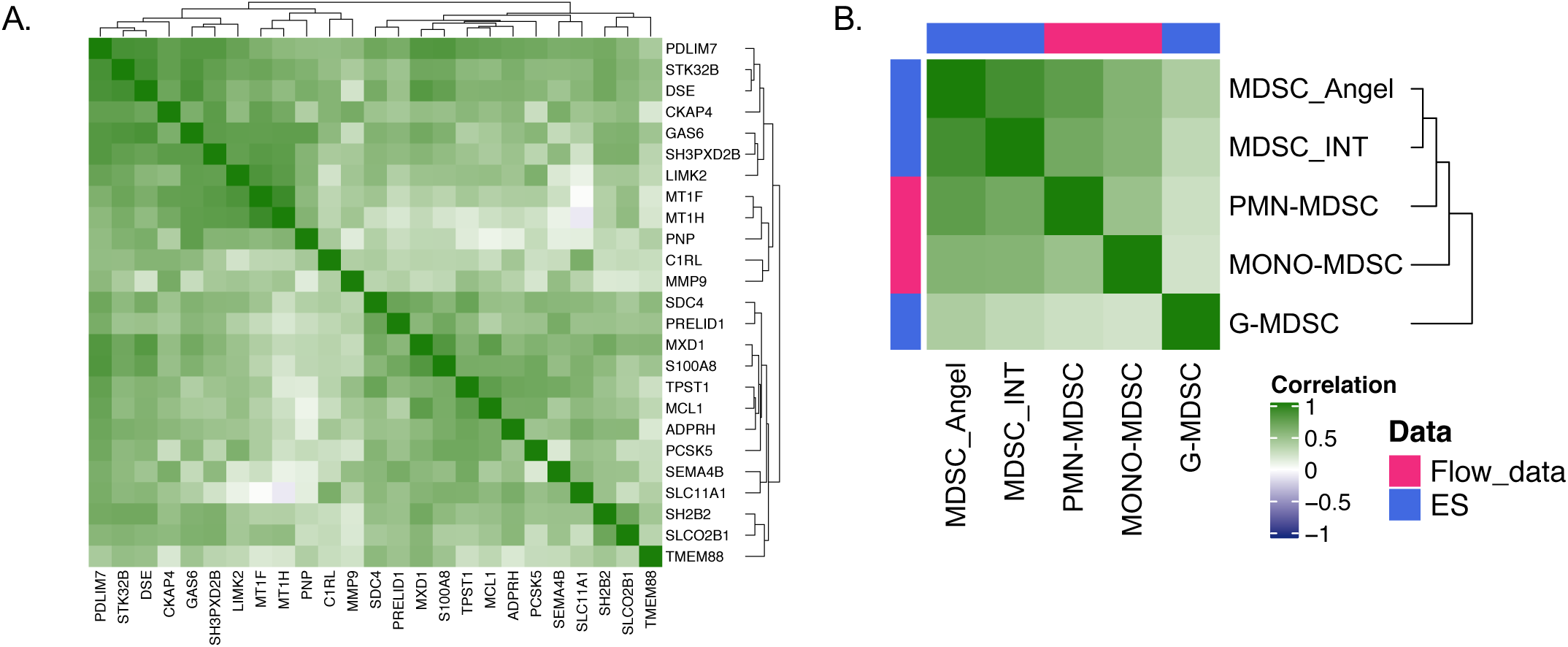
Pairwise Pearson correlation matrix between. **(A)** 25 genes of the top r2 (kmean1) of top 100 up-regulated genes from the data set obtained from human MDSC generated in vitro according to a model developed in our laboratory [Huber *et al*. JCI 2018]. **(B)** enrichment score of MDSC signature and percentage expression cell type by flow cytometry.

**Supplementary Figure 5:**
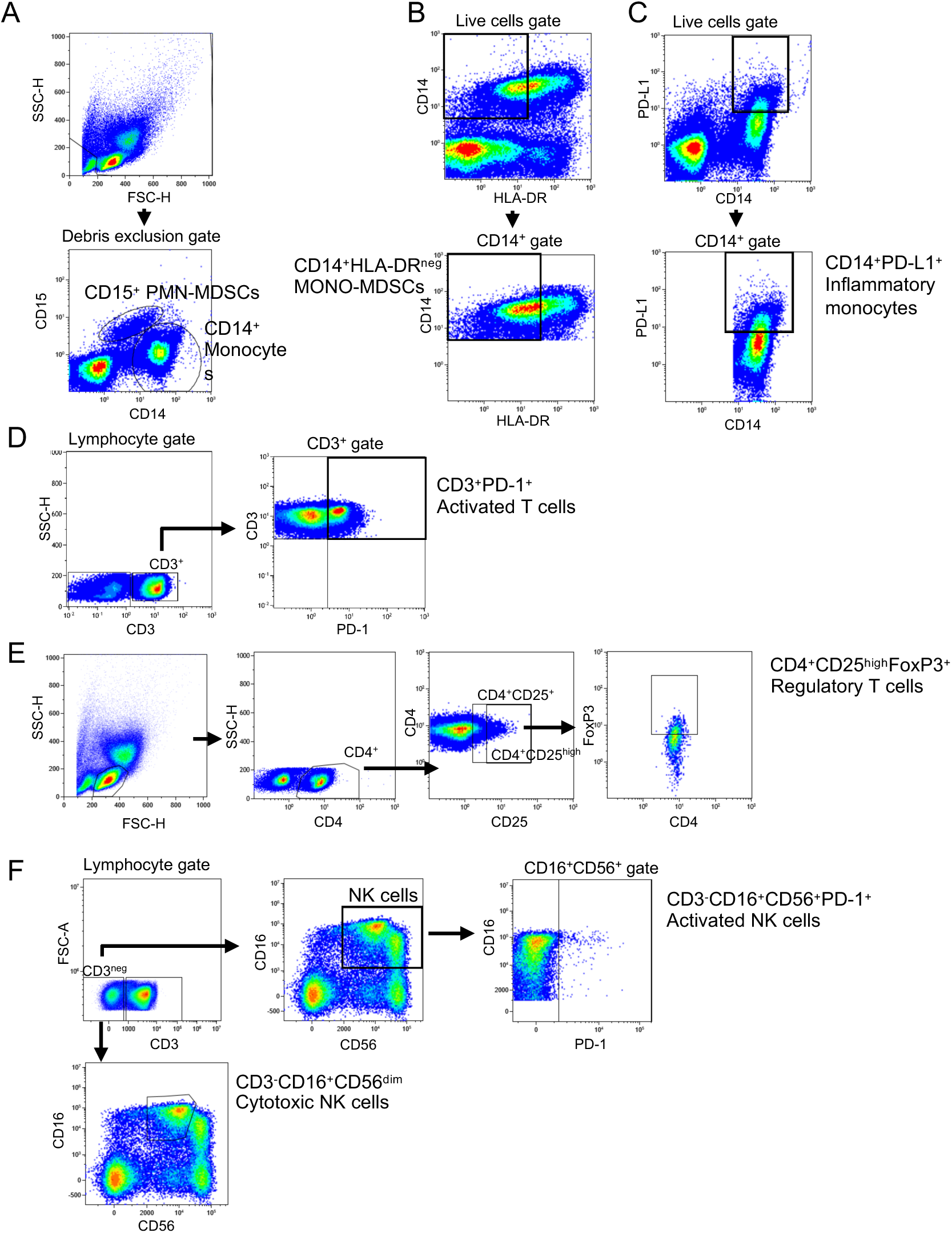
Flow cytometry gating strategies. Myeloid cell populations: CD15^+^ PMN-MDSCs, CD14^+^ monocytes, (A); CD14^+^HLA-DR^neg^ M-MDSCs (B); CD14^+^PD-L1^+^ inflammatory monocytes (C). T and NK cell populations: CD3^+^PD-1^+^ activated T cells (D); CD4^+^CD25^hi^ FoxP3^+^ regulatory T cells (E); CD16^+^CD56^+^PD-1^+^ activated NK cells and CD3^-^CD16^+^CD56^dim^ cytotoxic NK cells (F).

